# Wnt Signalling Orchestrates the Immune Response and Cardiac Damage During *Trypanosoma cruzi* Infection

**DOI:** 10.1101/2020.05.27.119107

**Authors:** Ximena Volpini, Laura Fernanda Ambrosio, Agustina Brajín, María Belen Brugo, María Pilar Aoki, Hector Walter Rivarola, Fernando Alfonso, Laura Fozzatti, Laura Cervi, Claudia Cristina Motran

## Abstract

Chagas’ cardiomyopathy is the consequence of a compromised electrical and mechanical cardiac function, with parasite persistence, unbalanced inflammation and pathological tissue remodelling, being intricately related to the myocardial aggression and the impaired function. Recent studies have shown that Wnt signalling pathways, which are important for developmental processes, play a critical role in the pathogenesis of cardiac and vascular diseases. In addition, we have reported that *Trypanosoma cruzi* infection activates Wnt signalling pathways in macrophages to promote their intracellular replication, with treatment of mice with IWP-L6 (an inhibitor of the O-acyl-transferase, PORCN, responsible for the post-translational modifications necessary for Wnt proteins secretion) being able to diminish parasitaemia and tissue parasitism. Therefore, Wnt signalling may contribute to the development of Chagas’ cardiomyopathy. In this work we have evaluated the effectiveness of Wnt secretion inhibition to control the parasite replication, modulate the adaptive immune response, and prevent the development of cardiac lesions in an experimental model of chronic Chagas disease. The IWP-L6 treatment, administered to *T. cruzi* infected BALB/c mice in a time window during the acute phase of the infection, was able to control the parasitaemia and heart parasitism together with the amelioration of the electrical, mechanical and histopathological cardiac alterations observed in chronically infected mice. Moreover, we demonstrated that during the acute phase of the infection Wnt signalling activation contributes to promote specific Th2-type immune response and to maintain the suppressive activity of Treg cells. Our data provide evidence that inhibition of Wnt signalling during the acute phase of *T. cruzi* infection controls the parasite replication, inhibits the development of parasite-prone and fibrosis-prone Th2-type immune response and prevents the development of cardiac lesions characteristics of chronic Chagas disease. Our study suggests that Wnt signalling pathway might be a potential target to prevent the development of *T. cruzi*-induced cardiomyopathy.

## Introduction

Chagas’ disease, caused by the infection with the protozoan parasite *Trypanosoma cruzi*, has an annual incidence of 30,000 cases in the region of the Americas affecting approximately 6 million people and causing an average of 12,000 deaths per year, along with the infection of 8,000 newborns during gestation annually. It is estimated that around 70 million people live in areas at risk of contracting the disease in the Americas; moreover, in the lest years, this infection is also becoming a worldwide health problem due to migration trends (1-3).

The disease presents two phases, acute phase, which last 6-8 weeks, and is followed by the chronic phase. The first one is characterized by high parasitaemia and tissue parasitism together with the activation of cells of innate immune components as dendritic cells (DCs), monocytes/macrophages (Mo), granulocytes and natural killer (NK) cells that instruct the development of a specific adaptive immune response. The first defence mechanism against the parasite is based on the release of IL-12 by infected Mo, which induces IFN-γ production from NK and T cells. In turn, IFN-γ activates Mo which control the intracellular parasite replication that occurs during acute infection, through the production of nitric oxide (NO), reactive nitrogen intermediates, reactive oxygen species (ROS) and kynurenines (4-8). In addition, TNF amplified microbicidal mechanisms of IFN-γ-activated Mo, thus contributing to host protection (9, 10). The cells of the innate immune system, through the production of cytokines and other effector molecules initiate the adaptive immune response. Together, CD4+ and CD8+ T cells as well as B-cells participate in controlling parasite multiplication and tissue invasion by cytokine secretion, cytotoxicity of infected cells and specific antibody production (11). Regarding the protective adaptive immune response, the simultaneous development of different T helper (Th) cell subsets and the outcome of *T. cruzi* infection could be defined as a battle between beneficial Th1-type response that fights against intracellular parasites, and Th2-type response (driven by alternatively activated Mo) that promotes parasite replication (12, 13). Additionally, IFN-γ- and TNF-secreting Th1 cells can orchestrate a deleterious response causing tissue destruction and fibrosis.(14) Thus, although required for parasite clearance, an increased Th1-type response with high levels of IFN-γ and TNF has been associated with the pathogenesis of chronic Chagas disease.(15-18).

Although *T. cruzi* can replicate within different nucleated cells; it shows particular tropism for striated cardiac myofibers, which explains why the heart is the most often affected organ in the chronic disease (19). Chagas’ cardiomyopathy is a slowly evolving inflammatory heart disease that develops in approximately one-third of *T. cruzi*-infected individuals and may lead to severe cardiac dilatation, congestive heart failure, and death (1). This cardiomyopathy is characterized by undetectable parasitemia, low tissue parasitism, focal areas of mononuclear infiltration, immune inflammation, necrosis and fibrosis of the myocardium that results in the appearance of abnormalities in the left ventricle (LV) function (1). Although the pathogenesis of Chagas’ heart disease is not completely understood, it is known that not only the parasite-mediated myocytolysis but also innate and adaptive immune mechanisms may contribute to heart damage through sustained inflammation and oxidative stress injury, leading to myofibrils disruption, myocyte necrosis, microvascular dysfunction, autonomic dysfunction, fibrosis and cardiac hypertrophy (1, 20).

Wnt proteins comprise a family of highly evolutionarily conserved secreted glycoproteins that bind transmembrane receptors of the Frizzled (Fzd) family and different co-receptors to trigger several signalling pathways that play central roles in organogenesis as well as in key fate decisions such as cell renewal, differentiation and apoptosis (21). Depending on the composition of the Wnt-Fzd-coreceptor complex and the cellular context, three Wnt signalling pathways have been characterized: the canonical Wnt/β-catenin pathway, the planar cell polarity pathway, and the noncanonical Wnt/Ca^+2^ pathway (22, 23). An imbalance in Wnt pathway activity has been implicated in the pathogenesis of different human diseases as cancer, degenerative disorders and inflammatory conditions (24). Wnt signalling pathways have a critical role in cardiac and vascular development and, therefore, is not surprising that they participate in cardiovascular diseases (25). Indeed, over the past years several studies have demonstrated that deregulated Wnt/β-catenin signalling pathway plays a key role in the induction of fibrosis, suggesting it as a novel therapeutic target in fibrotic disorders (26). Innate immune cells, as DCs and Mo, express a variety of Fzd receptors and Wnt proteins, and consequently they are susceptible to Wnt signalling regulation. Thus, several studies have highlighted the critical role of Wnt canonical pathway triggered on DCs and Mo in controlling the inflammatory responses to microbial infections (27, 28). Recently, we have reported that early after *T. cruzi* infection, the expression of some Wnt proteins, Fzd receptors as well as target genes of the Wnt/β-catenin pathway are induced with the Wnt canonical and non-canonical signalling pathways being activated in Mo (29). In addition, we have demonstrated that Wnt signalling has a critical role in modulating the innate immune response and the parasite replication during the acute phase of the infection. Treatment of C57BL/6 (B6) Mo with specific inhibitors of β-catenin transcriptional activity or with IWP-L6 (an inhibitor of the O-acyl-transferase, PORCN, responsible for the post-translational modifications necessary for Wnt proteins secretion) activates mechanisms that arm these cells to control intracellular parasite replication (29).

Moreover, *in vivo* treatment of B6 mice with IWP-L6 during a time window of the acute phase of the infection diminish the parasitemia and also reduce the parasite load in liver and heart during the chronic phase of the infection (29).

Considering that Wnt signalling pathways play relevant roles on two important aspects that exert influence on the development of Chagas’ cardiomyopathy, the balance between protective and pathogenic immune response and the orchestration of cardiac remodelling after injury (25), in this work we have evaluated the effectiveness of Wnt secretion inhibition to control the parasite replication, modulate the adaptive immune response, and prevent the development of cardiac lesions in an experimental model of chronic Chagas disease.

## MATERIALS AND METHODS

### Mice and Parasites

All animal experiments were approved by and conducted in accordance with guidelines of the Animal Care and Use Committee of the Facultad de Ciencias Químicas, Universidad Nacional de Córdoba (Approval Number HCD 743/18). BALB/c mice were obtained from Escuela de Veterinaria, Universidad Nacional de la Plata (La Plata, Argentina). All animals were housed in the Animal Facility of the Facultad de Ciencias Químicas, Universidad Nacional de Córdoba (OLAW Assurance number A5802-01). The Tulahuen strain of *T. cruzi* was used, which was maintained by weekly intraperitoneal (i.p.) inoculations in mice.

### IWP-L6 Treatment

Mice (6–8 weeks old) maintained under standard conditions were infected with 1,000 bloodstream *T. cruzi* trypomastigotes by the i.p. route. After 5, 8, 11, and 14 days post-infection (p.i.), mice were treated with IWP-L6 (7.5 mg/kg/dose) or vehicle by the i.p. route. IWP-L6 solution and blood parasite counts were assessed as described previously (29).

### Plasma Collection and Determination of CK and CK-MB

The blood was collect in heparinized tubes and centrifuged for 5 min at 2,300 r.p.m. to collect the plasma. The samples were derived to Biocon Laboratory (Córdoba, Argentina) for quantification of creatine kinase (CK) enzyme and CK-MB isoenzyme by UV kinetic method in a Dimension RXL Siemens analyzer.

### Cell Preparations and Culture

Spleen cell suspensions were obtained using tissue strainer and collected with PBS 2% FBS. Erythrocytes were lysed for 5 minutes in ammonium chloride-potassium phosphate buffer 0.87% (Sigma Aldrich). The number of cells was determined with Turk’s solution using Neubauer chamber. Total spleen leukocytes were resuspended in RPMI 1640 (Gibco, ThermoFisher Scientific) supplemented with 2 mM GlutaMAX (Gibco, ThermoFisher Scientific), 10% heat-inactivated fetal bovine serum (FBS, Natocor), and 50 µg/mL Gentamicin (Gibco, ThermoFisher Scientific). To evaluate specific immune response, splenic cells were cultured with *T. cruzi* lysate (10 μg/mL) at 2 × 10^6^ cells/mL for 72 h and the cytokines evaluated in the culture supernatant. Intracellular cytokines were detected after stimulating cells during 4 h with 10 μg/mL *T. cruzi* lysate in the presence of GolgiStop (BD Biosciences).

### Suppression Assay

Spleen cell suspensions from uninfected, control or IWP-L6-treated mice were obtained as described above and labeled with CD4-FITC and CD25-PECy7 antibodies for 20 min at 4°C. CD4+ CD25-splenic T cells (Teff) from uninfected mice and CD4+ CD25+ splenic T cells (Treg) from infected control or IWP-L6-treated mice were obtained by cell sorting with a FACSAria II (BD Biosciences). Teff cells were labeled by 5 min with CFSE (Invitrogen) and cocultured with Treg cells from infected control or IWP-L6-treated mice in a dish previously treated with anti-CD3 and anti-CD28 overnight at 4°C. Teff and Treg cells were cocultured at different ratios for 96 h. The suppressive capacity of Treg cells was assessed determining CFSE dilution by flow cytometry (FACSCanto II, BD Biosciences). The suppressive capacity of each Treg population was determined as the percentage of inhibition of Teff cell proliferation at Teff:Treg ratios 1:2; 1:1 and 2:1 relative to the maximal proliferation of Teff cells cultured alone (ratio 1:0).

### Flow Cytometry

Cells suspensions were incubated with fluorochrome labed-antibodies for superficial markers and intracellular cytokines detections as described previously (30). Different combinations of the following antibodies were used: CD3-APC, CD4-FITC, CD11b-PE, F480-PECy7, CD25-PECy7, Foxp3-PE, IFN-γ-PerCP5.5, TNF-PE, IL-4-APC, IL-5-PE, IL-10-PE and TGF-β-PECy7 (eBioscience). Samples were acquired in a FACSCanto II (BD Biosciences) and the data analyzed using FlowJO V10 software.

### Heart Homogenates

The hearts of mice were perfused with cold PBS and total homogenate suspensions were obtained using RIPA buffer (0.3 M NaCl, 20 mM Tris-HCl and pH 8.1% Sodium deoxycholate, 0.1% SDS,1% Triton X-100, 1 mM EDTA, 1 mM PMSF) supplemented with protease inhibitors (Protease Inhibitor Cocktail I, Tocris). Homogenates were centrifuged at 5000 g for 15 min at 4°C and supernatants were collected.

### Quantification of Released Cytokines

Cytokines were measured by capture ELISA according to the manufacturer’s instructions (eBiociences, ThermoFisher). Cytokine concentration in serum samples was expressed as pg/mL, while the cytokine levels in culture supernatants were represented as Index obtained by dividing the cytokine concentration in supernatant of *T. cruzi*-stimulated cultures by the cytokines concentration in supernatants of non-stimulated cultures (medium).

### Heart Parasitism and Histology

The apical section of the heart was used for determination of tissue parasitism as described previously (29). The rest of the organ was fixed in 10% buffered formalin and embedded in paraffin. Heart sections of 5 µm were examined after hematoxylin-eosin staining by light microscopy (Nikon Eclipse TE 2000 U).

### Electrocardiogram (ECG) and 2D-Echocardiography (2D-ECHO)

The ECG traces were obtained with standard potentials recorded at 50 mm/s with an amplitude of 1 mV/10 mm by the Contec ECG100G electrocardiograph. The alterations found were evaluated as previously described (7). The diastolic (DD) and systolic (DS) diameters of the left ventricle (LV) were studied by Logiq VDTM digital ultrasound. The LV shortening fraction (LVSF) was calculated as (DDVI-DSVI) / DDVI) x100, and the ejection volume (LVEF) was calculated from the same parameters by the Teichholz’s method. Uninfected BALB/c mice of the same age as infected experimental groups were included as controls of normal parameters.

### Antibody Assays

The quantification of IgG1 and IgG2a antibody isotypes against *T. cruzi* was carried out as previously described (31), using a *T. cruzi* lysate as antigen. Titrations were performed in duplicate and the end point was expressed as the serum dilution, with the optical density reading being twice that of the corresponding non infected sera.

### Statistical Analyses

Statistical analyses were performed using Student’s *t*-test, one- or two-way ANOVA followed by Bonferroni’s post-test as indicated by each figure. Each dot represents one animal. Statistical and graphs were performed by GraphPad Prism 8.0 software. The results were considered significantly different when *p* < 0.05.

## RESULTS

### Inhibition of Wnt Signalling Controls Parasite Load in Infected BALB/C Mice

During the infection with the Tulahuen strain of *T. cruzi*, BALB/c mice develop a cellular adaptive response biased to Th2-type, which is less efficient than Th1-type response to control this intracellular parasite infection (32). In this experimental model, the infection progresses chronically and is mainly characterized by intense cardiac inflammatory lesions that resemble in part the chronic cardiac pathology observed in human disease showing, several alterations of the cardiac electro-conduction and systolic function.

We have previously demonstrated that inhibition of Wnt signalling during the acute phase, improves the resistance to *T. cruzi* infection in B6 mice (29). To evaluate the effect of Wnt signalling inhibition on parasitaemia and heart parasitism in an animal model of chronic Chagas’disease, BALB/c mice were infected with a non-lethal doses of Tulahuen strain of *T. cruzi* and treated with IWP-L6 (IWP-L6 mice) or vehicle (Control mice) at days 5;8;11 and 14 p.i. Similar to that observed in B6 mice (29), IWP-L6 mice showed lower parasitaemia and heart parasite load than Control mice in the acute phase of the infection. (Figure 1A and B).

**Figure 1.**
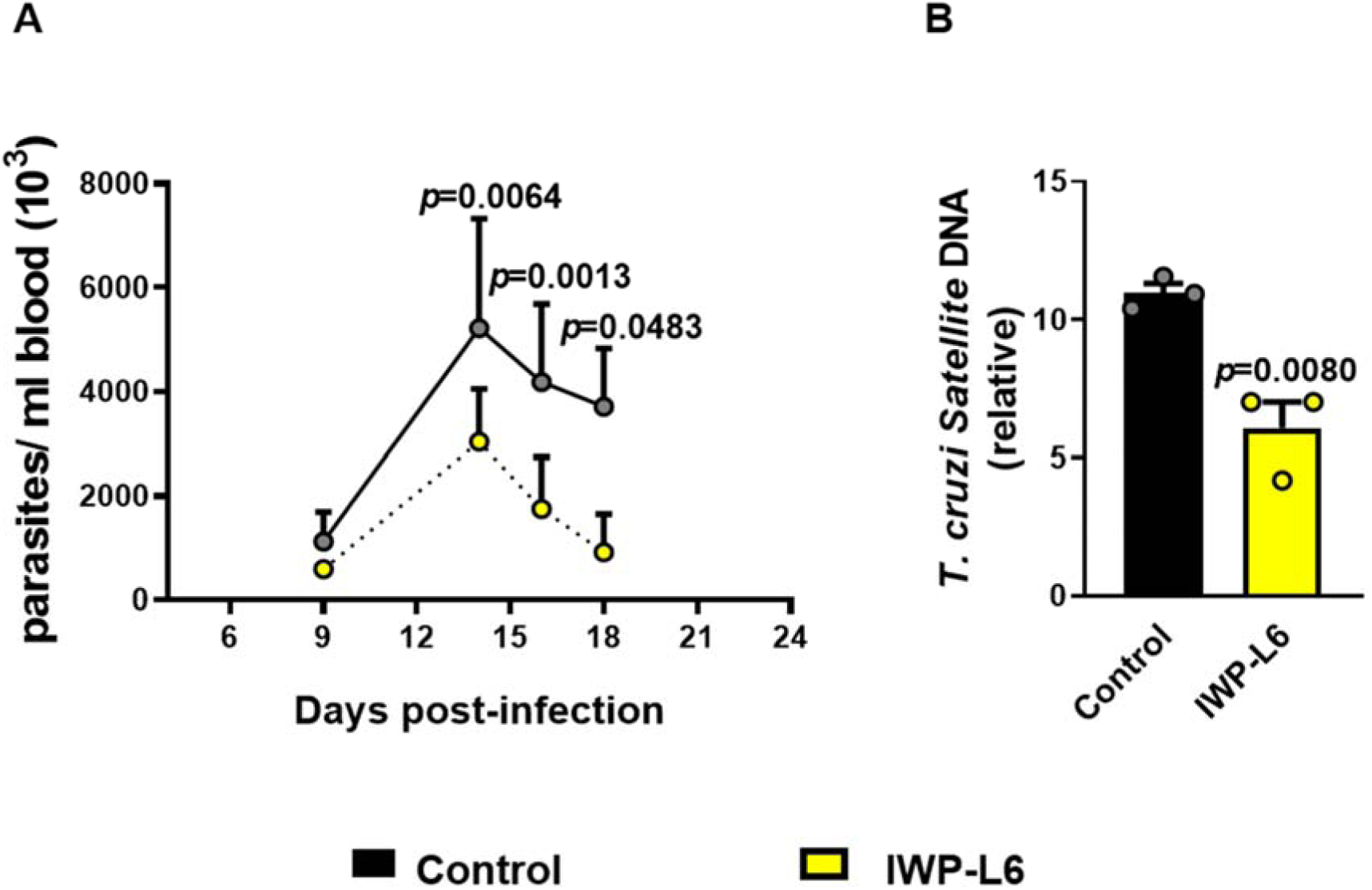
Effect of inhibition of Wnt signalling on parasitemia and heart parasite load. BALB/c mice infected with 1000 *T. cruzi* trypomastigotes were treated with IWP-L6 or vehicle on days 5, 8, 11, and 14 pi. **(A)** Parasitemia. Results are means ± SD of 8-9 animals/group and are representative of three independent experiments. **(B)** Relative amount of *T. cruzi* satellite DNA in heart from 18-day *T. cruzi-*infected IWP-L6 and Control mice. Murine GAPDH was used for normalization. Results are shown as means ± SD of 3 animals/group. Each symbol represents an individual mouse. Statistics were performed by two-ways ANOVA (A) or Student’ *t*-test (B). Experiment representative of 3.

### Inhibition of Wnt Signalling Promotes a Shift in Systemic Th1/Th2 Balance Toward Th1 Response at the Expense of Th2

To evaluate whether IWP-L6 treatment can modulate the typical Th2-type response mounted by BALB/c mice after *T. cruzi* infection (32), the levels of pro- and anti-inflammatory as well as Th1- and Th2-type distinctive cytokines were assessed in plasma at day 18 p.i. IWP-L6 mice showed significantly higher levels of IL-12 and TNF than Control mice, with the levels of IFN-γ, IL-4, IL-6 and IL-10 showing not significant differences between groups (Figure 2). Next, to assess the effect of IWP-L6 treatment on antigen specific response, splenic cells from IWP-L6 and Control mice obtained at 18 days p.i. were *in vitro* stimulated with an *T. cruzi* lysate antigen and the percent of Th1- and Th2-type cytokine producing cells and the profile of cytokines released to the culture supernatant was determined. Stimulated splenocytes from IWP-L6 mice secreted significantly higher amounts of IL-2 and IL-12, lower levels of IL-4 and similar levels of TNF, IFN-γ, IL-5, IL-10 and TGF-β compared with splenocytes from Control mice (Figure 3). Moreover, while no differences were observed neither in splenocytes total number nor in the frequency and absolute number of CD4+ T cells between both groups (Figure S1), significant lower percentage of CD3+ CD4+ IL-4 producing cells, and similar frequency of T cells producing IL-5 or the Th1-type cytokines (IFN-γ or TNF) were detected in stimulated spleen cells from IWP-L6 vs Control mice (Figure 4). Hence, the IWP-L6 treatment was able to modify the IL-4 to IFN-γ cytokine ratio calculated both as cytokine producing cells (Figure 5A) as well as *in vitro* supernatant concentration (Figure 5B). In accordance with having lower levels of IL-4, the IWP-L6 mice showed a significant reduction in the plasma levels of specific IgG1 antibodies (dependent on Th2 cells help (33)) in comparison with Control mice, without being affected the levels of specific IgG2a antibodies (Th1 help dependent(33)) (Figure 5C). Altogether, these data suggest that the *in vivo* inhibition of Wnt signalling promotes a shift in splenic Th1/Th2 balance toward Th1- at expense of Th2-type response.

**Figure 2.**
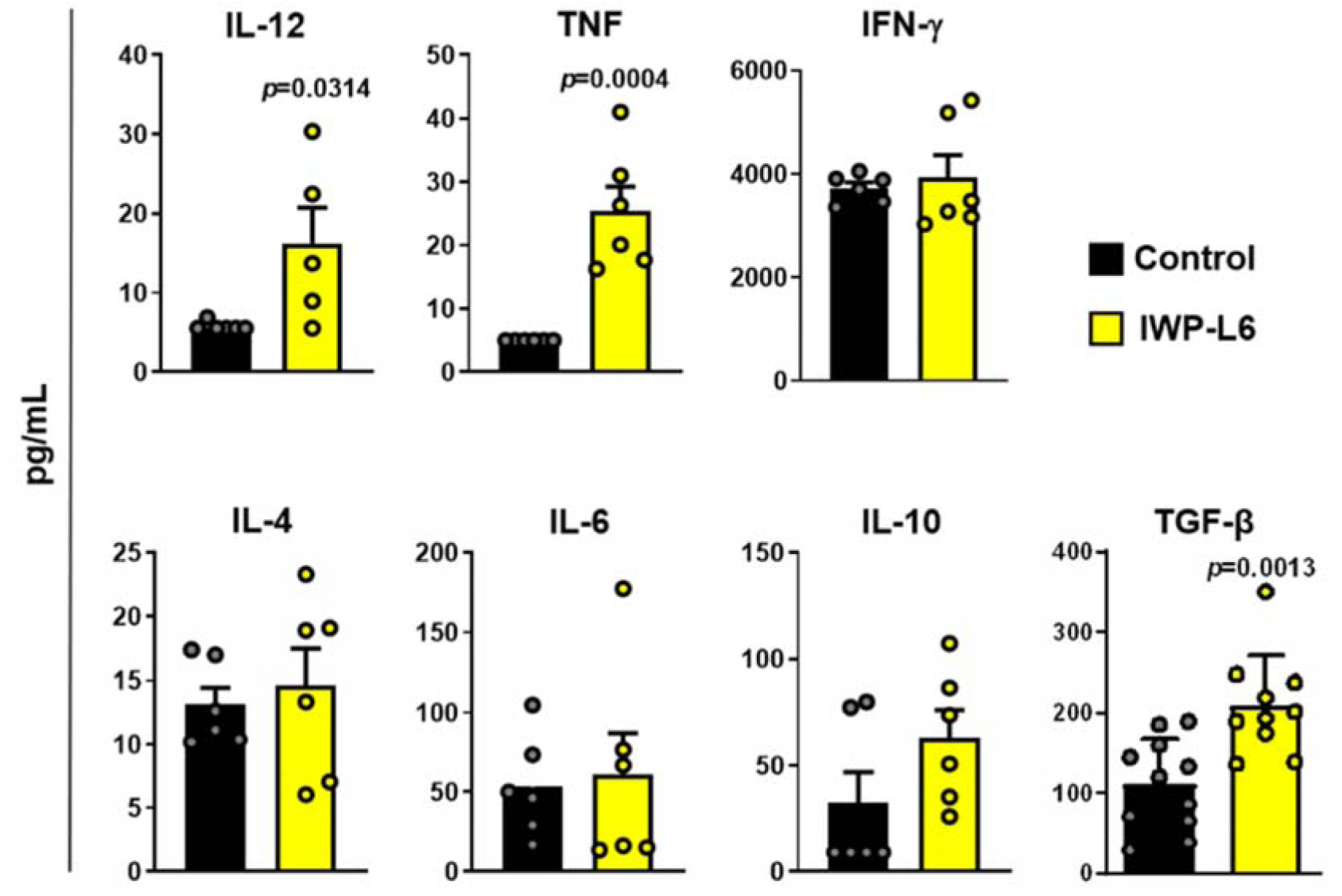
Plasma levels of pro- and anti-inflammatory cytokines in IWP-L6 and Control mice. BALB/c mice infected with 1000 *T. cruzi* trypomastigotes were treated with IWP-L6 or vehicle on days 5, 8, 11, and 14 pi. Plasma levels of cytokines were determined by ELISA on day 18 pi. Results are shown as means ± SD of 5-6 animals/group. Each symbol represents an individual mouse. Statistics were performed by Student’s *t*-test. Experiment representative of 3.

**Figure 3.**
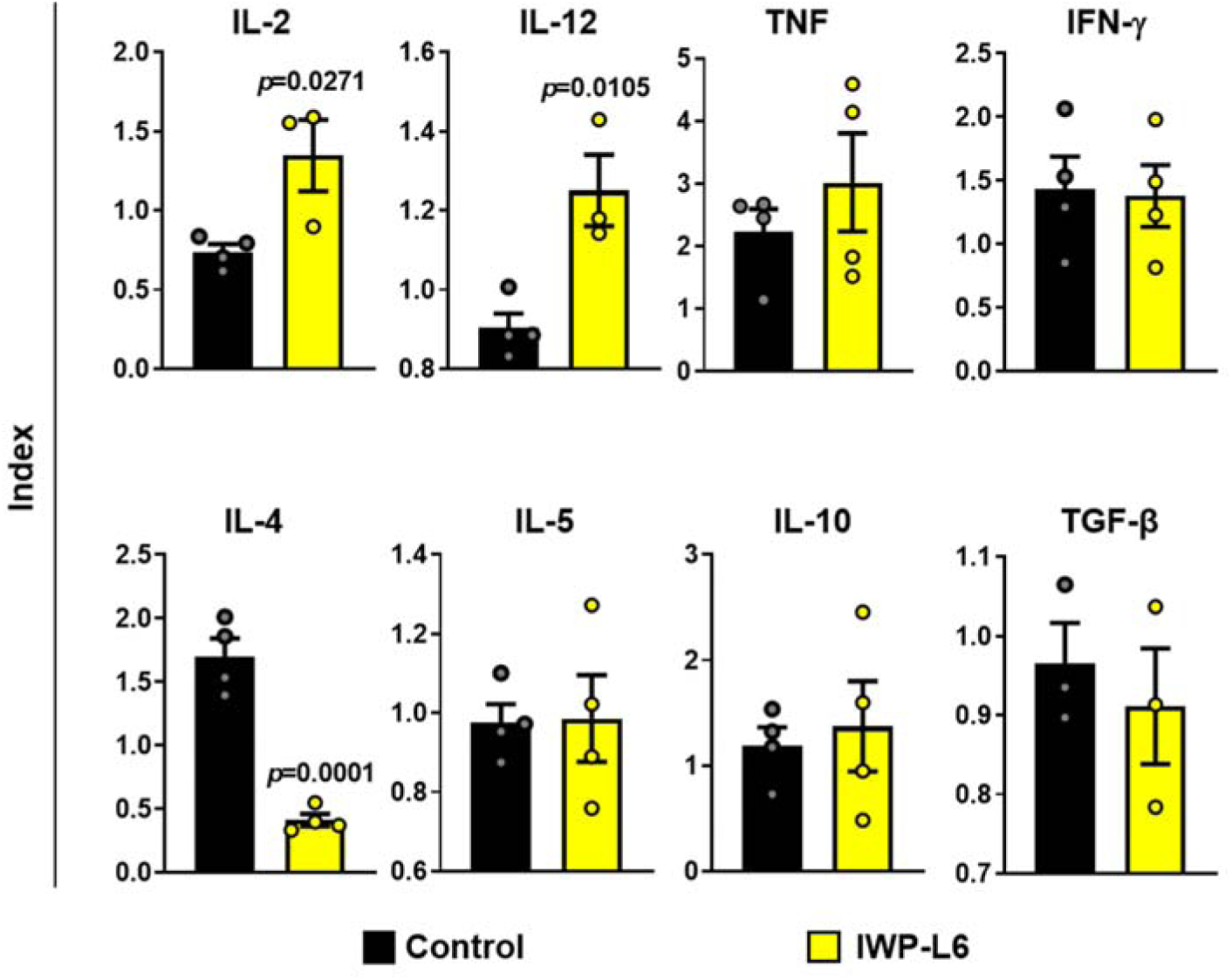
Cytokines secreted by *T. cruzi* stimulated splenocytes from IWP-L6 or Control mice. Splenocytes from IWP-L6 or Control mice obtained at day 18 pi were *in vitro* stimulated with *T. cruzi* antigens or medium alone for 72 h. The cytokines were quantified by ELISA in the culture supernatants and expressed as an Index obtained from the ratio between the concentration of cytokines released by splenocytes from each mouse cultured with *T. cruzi* antigens and cytokines released by splenocytes from the same animal cultured with medium. Results are shown as means ± SD of 3-4 animals/group. Each symbol represents an individual mouse. Statistics were performed using Student’s *t-*test. Experiment representative of 2.

**Figure 4.**
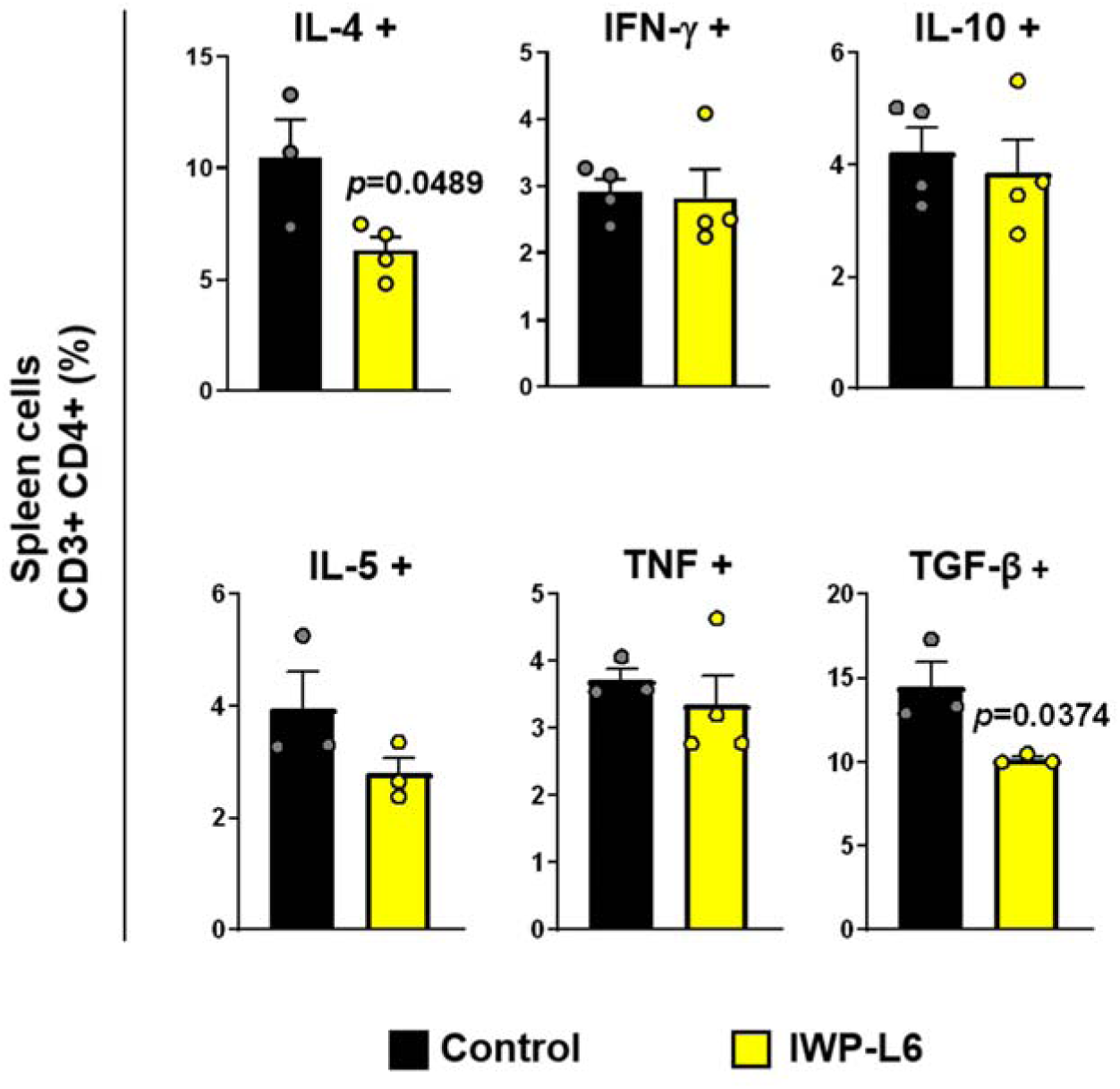
Effect of IWP-L6 treatment on cytokine production by splenic CD4+ T cells from *T. cruzi*-infected mice. Splenocytes from IWP-L6 or Control mice obtained at day 18 pi were *in vitro* stimulated with *T. cruzi* antigens or medium alone for 4 h in the presence of GolgiStop. Frequency of CD3+ CD4+ cells producing IL-4, IL-5, IFN-γ, TNF, IL-10 or TGF-β+ were determined by FACS. Results are shown as means ± SD of 3-4 animals/group. Each symbol represents an individual mouse. Statistics were performed by Student’s *t-*test. Experiment representative of 2.

**Figure 5.**
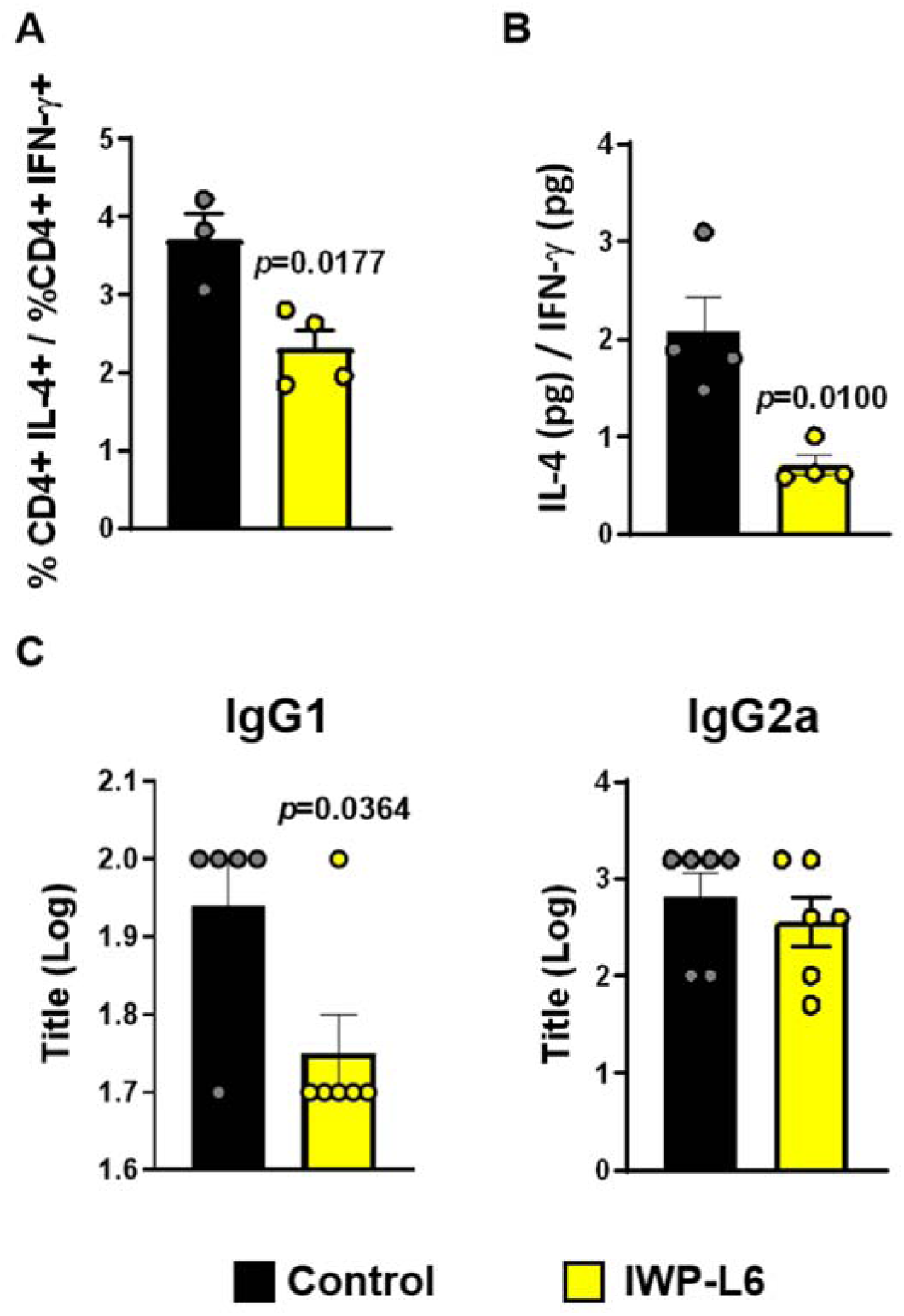
Effect of IWP-L6 treatment on Th2/Th1 balance. Splenocytes from IWP-L6 or Control mice obtained at day 18 pi were *in vitro* stimulated with *T. cruzi* antigens or medium alone, and the percentage of CD4+ T cells secreting IL-4 or IFN-γ and the levels of these cytokines secreted to the culture supernatant were evaluated as described in Fig. 3 and 4. **(A)** Ratio between the frequency of splenic CD4+ T cells from IWP-L6 or Control mice that produce IL-4 and IFN-γ. **(B)** Ratio between the concentration of IL-4 and IFN-γ determined by ELISA in culture supernatants of splenocytes from IWP-L6 or Control mice stimulated with *T. cruzi* antigens. **(A-B)** Results are means ± SD of 3-4 animals/group. One representative of two independent experiments is shown. **(C)** Plasma from IWP-L6 or Control mice obtained at 60 days p.i. were assayed for detection of IgG1 and IgG2a isotypes of antibodies against *T. cruzi* lisate by ELISA. Titrations were performed in duplicate, and the end point was expressed as the serum dilution with an optical density twice as high as those of the corresponding non-infected sera. Data are shown as the mean of the Log of antibody titles ± SD of 5-6 animals/group. Each symbol represents an individual mouse. Statistics were performed by Student’s *t-*test. Experiment representative of 3.

### Wnt Signalling is Important to Provide Functional Activity to Treg Cells From *T. cruzi*-Infected BALB/C Mice

The findings about the role of Wnt signalling during the differentiation and function of T regulatory (Treg) cells are still unclear and controversial. To evaluate the effects of Wnt signalling on Treg cells during *T. cruzi* infection, the absolute number and function of this splenic T cell population was assessed in IWP-L6 and Control mice at the peak of the parasitemia (18 days p.i.). Unlike B6 mice (34), *T. cruzi*-infected BALB/c mice expanded the population of Treg cells (CD4+CD25+Foxp3+) (Figure 6A) in parallel with the large expansion undergone by the T cell compartment (35). Therefore, the percentages of splenic Treg cells are similar in infected and uninfected mice (Figure 6B). Moreover, IWP-L6 and Control mice showed similar number and frequency of Treg cells in spleen (Figure 6A, B), suggesting that the inhibition of Wnt proteins secretion during a time window in the acute phase of the infection do not modify the Treg development or proliferation. Next, the functional properties of Treg cells from IWP-L6 and Control mice were comparatively analysed. To this aim splenic CD4+CD25+ (T_reg_) cells from IWP-L6 and Control mice were sorted and cultured with CD4+CD25- (T_eff_) cells sorted from uninfected BALB/c mice in the presence of anti-CD3 and anti-CD28 antibodies, with CD4+CD25+ cell population sorted from both groups of mice showing similar Foxp3 expression (percentages and mean fluorescence intensity, not shown). Suppression of T_eff_ cell proliferation by Treg cells from Control mice was observed at T_eff_/T_reg_ cell ratios of 1:2, 1:1 and 2:1, while Treg cells from IWP-L6 mice were unable to suppress T_eff_ cells proliferation at any rate assayed (Figure 6C), suggesting that during *T. cruzi* infection Wnt signalling contributes to maintain the suppressive activity of Treg cells. Furthermore, and in agreement with the lack of suppressive activity, splenic cells from IWP-L6 mice showed lower percentage of T cells producing the regulatory cytokine TGF-β than those from Control mice (Figure 4).

**Figure 6.**
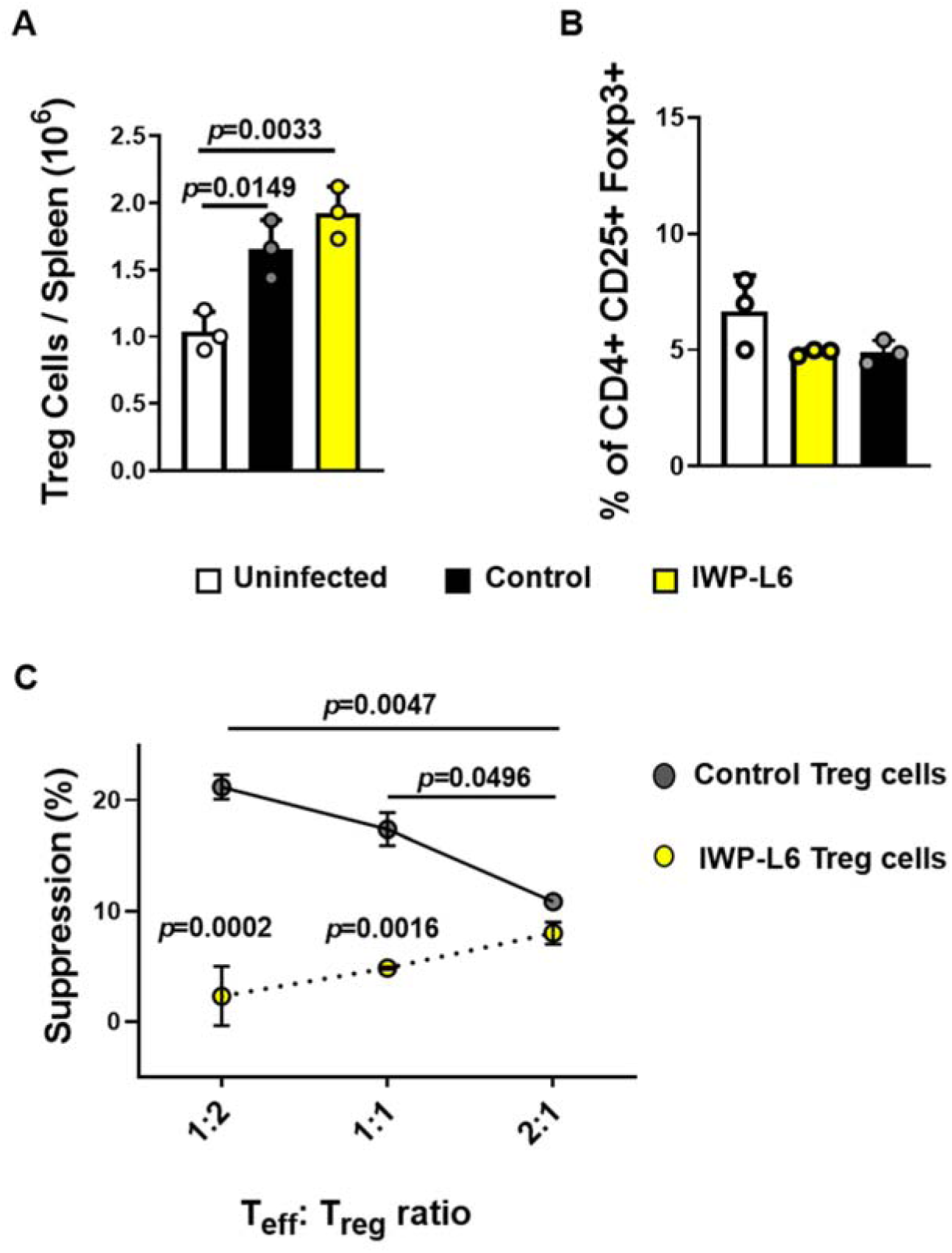
Effect of IWP-L6 treatment on Treg cells development and function. Absolute number **(A)** and percentages **(B)** of CD4+ CD25+ Foxp3+ cells in spleen from IWP-L6 or Control mice at 18 days pi. **(C)** Splenic CD4+ CD25-cells (T_eff_) sorted from uninfected BALB/c mice were CFSE labeled and cocultured with splenic CD4+ CD25+ cells (Treg) sorted from IWP-L6 or Control mice in plates coated with anti-CD3 and anti-CD28 for 96 h. The suppressive capacity of CD4+ CD25+ T cells was determined by FACS by evaluating CFSE dilution. Suppressive capacity of Treg cells was shown as the percentage of inhibition of Teff proliferation at T_eff_:Treg ratios of 1:2; 1:1 and 2:1 relative to the maximal proliferation of Teff cells cultured alone (ratio 1:0). Values are shown as means ± SD of 3 animals/group. All data are from one experiment representative of three in total. Statistics were performed by one-way (A) or two-ways ANOVA (B).

### Wnt Signalling Contributes to Impair Cardiac Function During *T. cruzi* Infection

The Chagas’ heart disease established after several years of primo infection is the consequence of compromised electrical and mechanical cardiac function induced by fibrosis and lymphocytic infiltration, which lead to tissue remodelling. To evaluate the role of Wnt signalling in the development of the cardiomyopathy characteristic of chronic *T. cruzi* infection, the effect of blocking Wnt signalling with IWP-L6 given in a small window of time during the early phase of the infection was assessed by determining the levels of serum markers of myocardial damage at the acute phase and the electrical, mechanical and histopathological cardiac alterations at the chronic phase of the infection. Figure 7A shows that the levels of CK a CKMB determined at 18 days p.i. were significantly increased in Control vs uninfected mice, with IWP-L6 mice showing significant lower CKMB levels than Control mice. Furthermore, the histopathological evaluation revealed extensive perivascular inflammatory focus and severe fibrosis in cardiac tissue of Control mice, while only remote inflammatory focuses were observed in IWP-L6 mice (Figure 7B). These findings were consistent with more severe electrical and mechanical abnormalities observed in Control vs IWP-L6 mice (at 180 days p.i.) (Figure 8). Indeed, the analysis of ECG traces (Figure 8A), demonstrated that whereas ECG from Control mice presented increased QT and PR intervals, the pathognomonic electrophysiological heart alteration observed in *T. cruzi-*infected BALB/c mice (36), ECG from IWP-L6 mice showed normal values of QT, indicating a protective cardiac role of IWP-L6 treatment (Figure 8B). Besides, Chagas’ heart disease is characterized by the dilatation of the LV and its consequent dysfunction (37). In agreement with the other results, two-dimensions echocardiography (2D-ECHO) study revealed that untreated infected mice exhibited a marked reduction in the LV percent fractional shortening (LVFS) and LV ejection function (LVEF), with the reduction of these both parameters being attenuated by treatment with IWP-L6 (Figure 8C). Moreover, LVEF and LVFS values of IWP-L6 mice showed not significant differences with those observed in uninfected mice (Figure 8C). Collectively, these data suggest that during *T. cruzi* infection the Wnt signalling system plays a critical role in orchestrating the cardiac damage.

**Figure 7.**
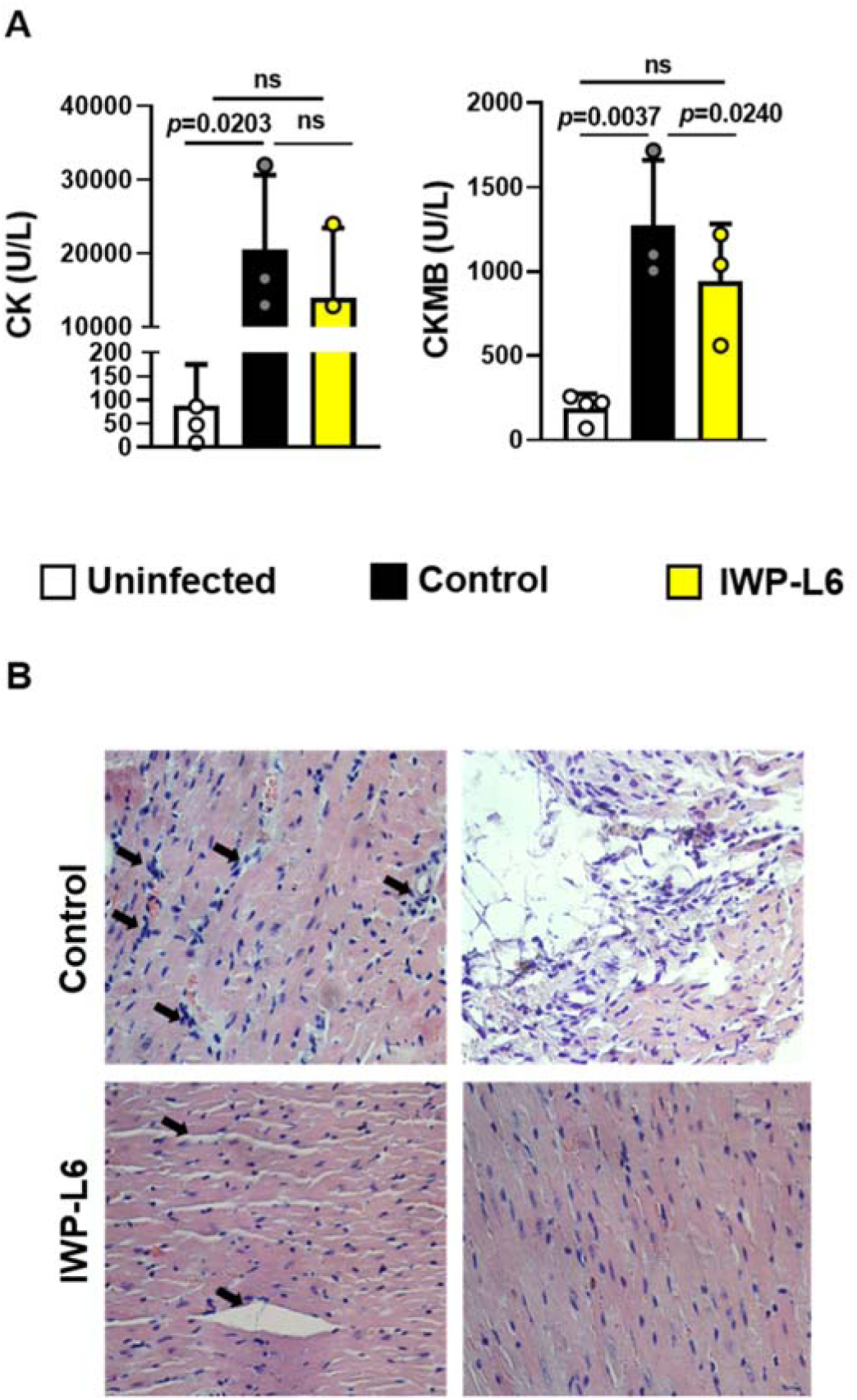
IWP-L6 treatment impairs *T. cruzi*-induced cardiac damage. **(A)** Plasma levels of CK and CK-MB at 18 days pi. Values are shown as means ± SD of 3 animals/group. Each symbol represents an individual mouse. Statistics were performed by one-way ANOVA. Experiment representative of 3. **(B)** Representative histological sections of heart from IWP-L6 or Control mice obtained at 180 days p.i. were stained with H&E and observed under light microscope (magnifications 1000X). Left panels indicate inflammatory infiltrate located close to the blood vessels (arrows) while right panels show parenchymal fibrosis. Graph is representative of one microscopy image of 5-7 animals/group of two independent experiments.

**Figure 8.**
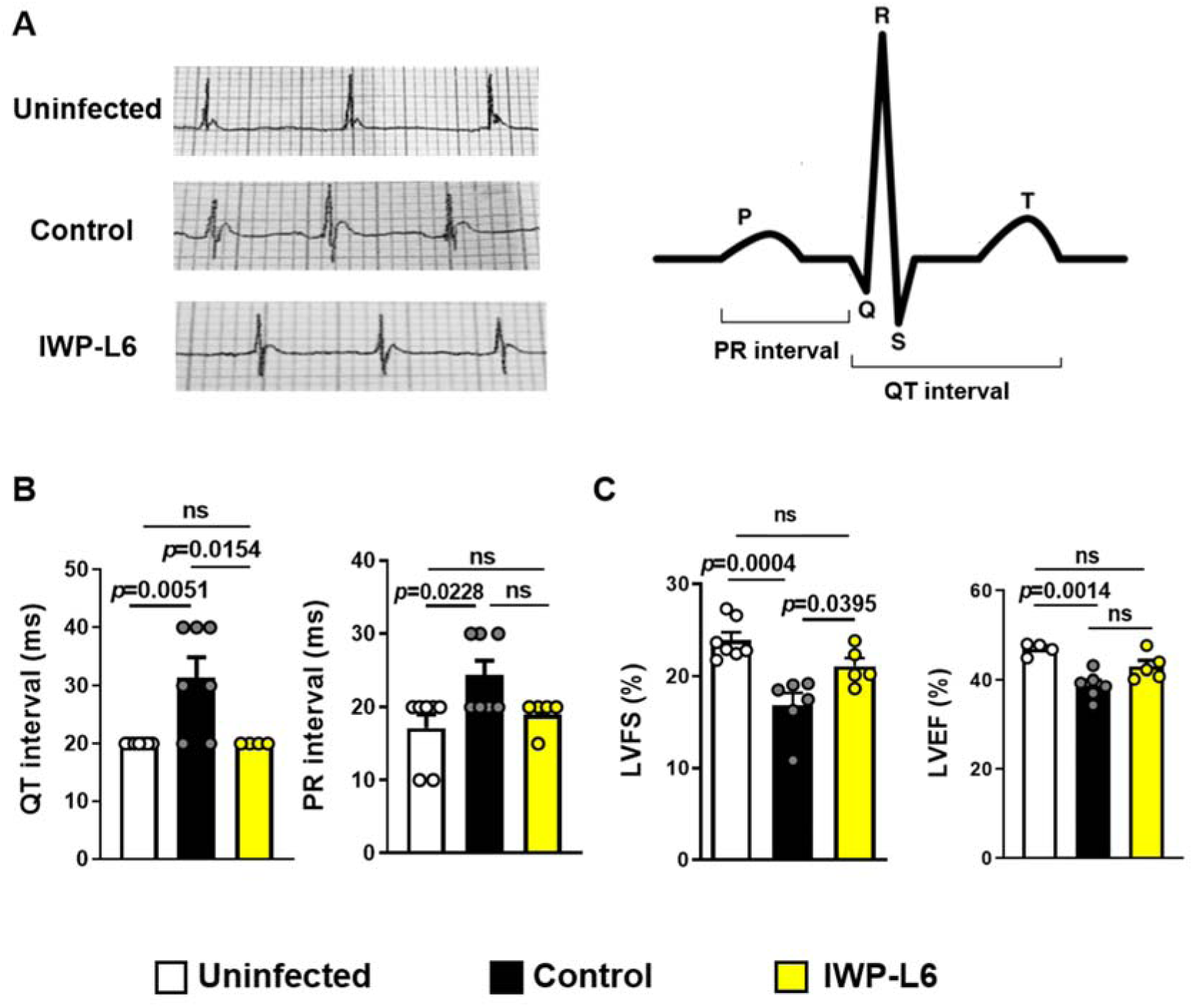
IWP-L6 treatment prevents the development of chronic Chagas’ disease-associated functional pathology. BALB/c mice infected with 1000 *T. cruzi* trypomastigotes were treated with IWP-L6 or vehicle on days 5, 8, 11, and 14 pi. ECG and ECHO heart function of uninfected, Control and IWP-L6 mice at 180 days p.i. **(A)** Left panel: Representative tracings for each group. Right panel: schematic representation of QT and PR intervals. **(B)** Values of QT and PR intervals of ECG from uninfected, Control and IWP-L6 mice. **(C)** Values of left ventricular fractional shortening (LVFS) and left ventricular ejection fraction (LVEF) of ECHO from uninfected, Control and IWP-L6 mice. Results are shown as means ± SD of 5-7 animals/group. Each symbol represents an individual mouse. Statistics were performed by one-way ANOVA. Experiment representative of 2.

## DISCUSSION

The therapies currently used for Chagas’ disease treatment are focused on parasitological cure and the control of the symptoms, with Benznidazole and Nifurtimox being the only drugs with proven trypanocidal efficacy (37). However, although having good and reasonably efficacy administered during the acute phase, the results obtained during the chronic disease are not conclusive. Both treatments have several side effects and current data indicate that in established cardiac disease, the treatment unlikely change clinical outcomes (38, 39). Thus, drugs inefficiency, the lack of an effective vaccine able to prevent the establishment of the infection and the persistence of insect vector in endemic areas, are all factors that increase the prevalence of the disease (40).

The scarcity of parasites in the heart of chronically infected individuals (41) has triggered a discussion regarding the etiology of cardiomyopathy. Nevertheless, nowadays it is well accepted that cardiac parasite persistence positive correlates with myocarditis degree (42). Because both host innate and adaptive immune responses are critical factors for inducing parasite clearance and tissue damage or protection, an effective immunomodulatory treatment should be able to control parasite replication and mount an effective balance between the host protective immune response and disease tolerance. On the other hand, it is important to take into account that not only immune system cells and their products but also many other signalling systems and complex cellular interactions between recruited and cardiac cells (myocytes, fibroblasts, endothelial cells, smooth muscle cells, epicardial cells) can participate in the damage and repair processes that occur after the parasite- or inflammation-induced injury. Thus, during *T. cruzi* infection, the Wnt signalling system would not only be involved in modulating Mo phenotype and function, as we have previously demonstrated (29), but could also play a role in orchestrating the cardiac damage that follows a parasite infection as it has been shown in ischemia or hypertrophy (reviewed by Deb, A (43)). Here, we show evidences that inhibition of Wnt was able to control the parasite load, modulate the parasite-prone and fibrosis-prone Th2-type immune response and prevent the development of cardiac lesions in an experimental model of chronic Chagas disease.

In contrast to B6 mice that mount an specific Th1-type response characterized by IL-12, IL-18 and IFN-γ as well as NO-producing M1-like Mo that control parasite replication at the early phases of the infection (44), *T. cruzi-*infected BALB/c mice develop a Th2-type cytokine environment that induce alternatively activated Mo (M2-like) showing high arginase-1 activity that favours the establishment of the parasite in the host (45). Considering that inhibition of Wnt signalling pathways using IWP-L6 before infection arm Mo to fight against *T. cruzi*, by inducing a subtype of Mo showing enhanced production of pro-inflammatory cytokines and lower arginase-1 activity (29), we speculate that this treatment contributes to shift the characteristic Th2-type immune response developed by BALB/c mice to a Th1-type. However, we found that despite being able to control parasite load, infected BALB/c mice that were treated with IWP-L6 did not present a significant increase in the Th1 parasite-specific immune response. Instead, they showed a pronounced decrease in specific Th2 response, pro-fibrotic and parasite prone, which resulted protective against the infection. Although the role for Wnt signalling pathways in regulation of T cell development has been extensively demonstrated, the role of T cell factor (TCF), an effector transcription factor (TF) of the canonical Wnt pathway, in Th cell differentiation and function has been recently addressed (46). Thus, it has been reported that TCF-1 act as a cofactor of β-catenin to induce in TCR-activated T cells the expression of GATA-3 TF, required for Th2 differentiation, in an IL4-independent fashion (47, 48). Regarding to Th1 differentiation, the percentage of IFN-γ-producing T cells is increased in TCF-1-deficient mice with TCF-1 mRNA being downregulated in naïve CD4+ cells that are differentiated into Th1 profile (47). In addition, it has been reported that TCF-1 is part of a transcriptional complex that impairs T-bet expression, limiting its capacity to induce IFN-γ expression (49), although in the primary response to virus infection the lack of TCF-1-β-catenin binding do not modify the capacity of CD4+ T cells to express IFN-γ or T-bet (50). Therefore, our results are consistent with the notion that canonical Wnt pathway activation promotes Th2 CD4+ T cell differentiation and represses Th1 phenotype, although in the present work Th1 profile expansion was not observed after Wnt signalling inhibition. Nevertheless, the fact that the levels of the Th1-type pro-inflammatory cytokines TNF and IL-12 but not NO (not shown) were higher in plasma from IWP-L6 vs Control mice, together with the higher capacity of splenocytes from IWP-L6 mice to secrete IL-12 after *in vitro* antigen-specific stimulation, suggest that IWP-L6 treatment promotes *in vivo* the activation of Mo to an M1-like phenotype that secretes pro-inflammatory cytokines but not produces NO and are more efficient to control the parasite load, as we demonstrated in B6 mice (29).

We and others have shown that during the early phase of *T. cruzi* infection, as in other parasitic diseases, the limitation of Treg cells allow the emergence of the protective T cell-dependent response (30, 34, 51, 52). However, reduction in Treg frequency can result in a massive accumulation of effector immune cells and immunopathology, as is observed in *T. cruzi*-infected B6 mice (34, 53). In contrast to that observed in infected B6 mice which are unable to induce Treg cells (34, 53), here we demonstrated that infected BALB/c mice expand both effector and Treg cells compartments during the acute phase of the infection, maintaining the frequency of Treg cells within normal limits. Thus, *T. cruzi* infected BALB/c mice show limited Th1-type protective response with a predominance of Th2-type response, but they are less susceptible than B6 mice to acute immunopathology (32). The development of Treg cells is orchestrated by a series of signalling pathways. TCF-1 and LEF-1, which act downstream Wnt/β-catenin pathway as positive mediators of β-catenin transcriptional activity, are among the TF that work in synergy with Foxp3 to activate the expression of most of the Treg transcriptional program (54). Different studies have shown that strong or persistent Wnt stimulation inhibit Treg cells suppressive function (55-57). However, forced expression of a stabilized form of β-catenin enhanced Treg cell survival and conferred better protective capacity to Treg cells in a colitis model (58). Moreover, recently two different groups have demonstrated that TCF-1 and LEF-1 expression are critical for sustaining Treg suppressive functions to prevent from autoimmunity (59, 60). Hence, the role of Wnt signalling pathway on Treg cells is still controversial. In our study, Treg cells development or proliferation was not modified by IWP-L6 treatment. However, CD4+ CD25+ splenic cells from IWP-L6-mice did not inhibited the proliferation of T_eff_ cells as did those from Control mice. This observation is in agreement with the lower percentages of splenic CD4+ T cells producing TGF-β, a cytokine used by Treg cells to control effector T cells (61). Taken together the results, we found that Treg cells function was impaired after Wnt signalling inhibition with IWP-L6. Interestingly, in agreement with our results, inhibition of Wnt/β-catenin signalling in a model of lung fibrosis, restricts Treg cells suppressive capacity, the production of TGF-β and the Th2 polarization (62). Since several reports have demonstrated that Wnt/β-catenin signalling pathway negatively modulates the function of Treg cells (55-57), we expected that Wnt signalling blockade enhances the suppressive ability of Treg cells. However, several differences may help to explain the apparently opposing results obtained by using an *in vivo* model of parasitic infection and those from other groups using different experimental settings. While genetic or pharmacological gain- or loss-of-function approaches target only the Wnt/β-catenin signalling pathway, we have demonstrated that *T. cruzi* infection induces canonical but also Wnt/Ca++ activation, and therefore both pathways are targeted by IWP-L6 treatment (29) in the context of this infection. Besides, additional research is required to define Wnt-Fzd receptors combinations relevant for Treg cells function regulation, and how inflammatory signals can modulate Wnt signalling pathways *in vivo*. Because the inflammatory microenvironment of cancer (tolerogenic) or autoimmune disease (highly inflammatory) are very different from that in the context of parasite infection, the type and levels of Wnt proteins and Fzd receptors that are up- or down-regulated in Treg cells could be also different. Also, since Wnt/β-catenin signalling in DCs promotes the expression of anti-inflammatory mediators such as TGF-β and the induction of Treg cells (63-65), it is important to consider that the blocking of Wnt signalling by IWP-L6 treatment also affects DCs.

Chagas’ cardiomyopathy is the leading cause of mortality of infected patients, due to compromised electrical and mechanical cardiac function induced by lymphocytic infiltration, cardiomyocyte hypertrophy and prominent fibrosis (1, 66). Accordingly, the murine *T. cruzi* infection model shows alterations in the cardiac electrical conduction system characterized by an atrioventricular blockage, identified by irregular and increased QT intervals, cardiac arrhythmias and sinus bradycardia (36). Different studies have demonstrated that during pathologic remodelling of the heart, several foetal genes, including components of Wnt signalling, become reactivated (67). Thus, changes in Wnt expression or aberrant Wnt/β-catenin signalling have been reported in different cardiac diseases as cardiac hypertrophy, ischemic cardiac injury and arrhythmogenic cardiomyopathy. Intervention at the level of either inhibition of Wnt proteins secretion, Fdz, the β-catenin destruction complex, or β-catenin-mediated gene transcription after ischemic cardiac injury have all shown beneficial effects on infarct size, fibrosis, cardiac function and LV remodelling (reviewed by Foulquier et al. (25)). Moreover, β-catenin loss-of-function mutations in myocytes or overexpression of GSK-3β were associated with enhanced cardiac function in cardiac hypertrophy models (68-71). Therefore, activation of Wnt/β-catenin signalling pathway plays a dominant role in the regulation of cardiac fibrosis and pathological remodelling after cardiac injury (72, 73). On the other hand, non-canonical Wnt signalling pathways are most likely involved in orchestrating the inflammatory processes in ischemic heart (74). Furthermore, several reports have implicated that PORCN inhibitors, identified as novel antitumor drugs and applied in ongoing clinical trials, are potential drugs for treatment of myocardial infarction acting through mechanisms that include reducing cardiomyocyte death, increasing angiogenesis, suppressing fibrosis and stimulating cardiac regeneration (75-77). In agreement with these results, we have found that the temporal treatment with IWP-L6, a PORCN inhibitor, preserved electric cardiac function, reduced cardiac damage and myocarditis and prevented the thinning of the LV wall, which retain their ejection capacity. As it has been reported that during *T. cruzi* infection of cardiomyocytes the activation of Ca^++^/Calcineurin/NFAT axis promotes the production of inflammatory mediators that induces cardiac damage (78), it is possible that during *T. cruzi* infection, the activation of both Wnt canonical and non-canonical signalling pathways are contributing to the development of cardiac pathology. Hence, parasite-induced myocytolysis and myofibrillar degeneration, intense cell infiltration and exacerbated inflammatory response together with Wnt signalling pathways activation would be involved in the damage, subsequent fibrosis and cardiac function alterations during *T. cruzi* infection (1, 25). All these variables are related and depend on each other, because the cardiac damage and inflammation are linked to the expression of Wnt proteins and Fdz receptors and the activation of Wnt signalling on immune and non-immune cells from “cardiac interactome” (29, 43, 79). In addition, heart parasitism which is strongly related to cardiac damage and inflammation (80), and Th2-type response which is linked to fibrosis (81), were significantly lower in IWP-L6 than in Control mice. Therefore, to discuss the mechanisms that operate to protect cardiac architecture and function after IWP-L6 treatment it is important to keep in mind that this treatment, through the modulation of innate and adaptive immune response, is controlling the parasite load.

Taking into account that the development of Chagas cardiomyopathy is associated with cardiac parasite persistence and an excessive inflammatory response and fibrosis, our finding suggests that the temporal treatment with the PORCN inhibitors limits parasite replication, diminishes Th2-type pro-fibrotic adaptive immunity, and inhibits the production of inflammatory mediators, improving the functionality of infected heart.

## Acknowledgments

This work was supported by grants from Consejo Nacional de Investigaciones Científicas y Técnicas from Argentina (CONICET), Agencia Nacional de Promoción Científica y Técnica (PICT 2015-2488, PICT 2016-0415) and Secretaría de Ciencia y Técnica, Universidad Nacional de Córdoba (grants to CCM). The funders had no role in study design, data collection and interpretation, or the decision to submit the work for publication.

XV and LFA thank Consejo Nacional de Investigaciones Científicas y Técnicas from Argentina for the fellowships granted. AB and MBB thank Agencia Nacional de Promoción Científica y Técnica for the fellowships granted. CCM, MPA, LF and LC are members of the Scientific Career of Consejo Nacional de Investigaciones Científicas y Técnicas from Argentina. We thank S. Miró, F. Navarro, D. Lutti, V. Blanco, C. Noriega, A. Romero, G. Furlan, L. Gatica, MP Abadie and P. Crespo for their excellent technical assistance. We are grateful for the assistant of E. Acosta Rodriguez, C. Araujo Furlan and F. Canale for help in suppression assay.

XV designed and performed most of the experiments, analyzed data, and wrote/commented on the manuscript. LA, AB, CR, MBB, MPA, HWR, FA performed experiments and commented on the manuscript. LF, MPA and LC participated in data analysis and commented on the manuscript. CCM supervised the research, designed experiments, wrote the manuscript, and provided funding.

