## Supplementary Material for "Wnt Signalling Orchestrates the Immune Response and Cardiac Damage During *Trypanosoma cruzi* Infection"

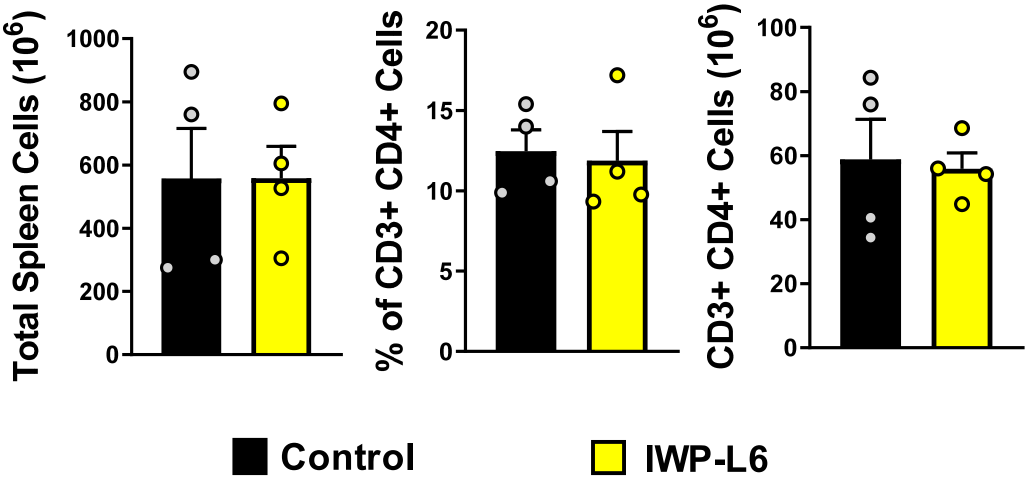


**Supplementary Figure 1.** Splenocytes from IWP-L6 or Control mice obtained at day 18 pi were counted (left panel) and stained with anti-CD3 and -CD4 for the determination of the percentage of CD3+ CD4+ cells/spleen (middle panel) and the absolute number of CD3+ CD4+ cells/spleen (right panel) by FACS. Results are shown as means ± SD of 4 animals/group. Each symbol represents an individual mouse. Statistics were performed by Student's *t*-test. Experiment representative of 3
